# A fluorescent assay for cryptic transcription in *Saccharomyces cerevisiae* reveals novel insights into factors that stabilize chromatin structure on newly replicated chromatin

**DOI:** 10.1101/2023.07.11.548543

**Authors:** Ellia Gao, Stephanie Jung, LeAnn J. Howe

**Affiliations:** Department of Biochemistry and Molecular Biology and Life Sciences Institute, The University of British Columbia, 2350 Health Sciences Mall, Vancouver, B.C., Canada, V6T 1Z3

**Keywords:** cryptic transcription, chromatin, DNA replication

## Abstract

The disruption of chromatin structure can result in transcription initiating from cryptic promoters. A well-characterized, chromatin-destabilizing stress is the passage of RNA polymerase, and numerous factors function to stabilize chromatin on transcribed genes, suppressing cryptic transcription from sites within gene bodies. DNA replication is also inherently disruptive to chromatin, and multiple replication-coupled histone chaperones suppress cryptic transcription. However, these factors also have documented roles in transcription, and thus whether DNA replication per se can activate cryptic promoters has not been directly examined. In this study, we tested the hypothesis that, in the absence of chromatin-stabilizing factors, DNA replication can promote cryptic transcription in *S. cerevisiae*. Using a novel fluorescent reporter assay, we show that multiple factors, including Asf1, Rtt106, Spt6, and Spt16, suppress transcription from a cryptic promoter, but are entirely or partially dispensable in G1-arrested cells, suggesting a requirement for DNA replication in chromatin disruption. Additionally, for the first time, we demonstrate modest cryptic transcription following the depletion of Rlf2/Cac1, a CAF-1 chromatin assembly complex component. Collectively, these results suggest that transcription fidelity is dependent on numerous factors that function to assemble chromatin on nascent DNA.

## Introduction

Eukaryotic DNA is packaged into chromatin, a nucleoprotein structure in which DNA is wrapped around octamers of core histone proteins. In addition to facilitating genome compaction through the neutralization of DNA charge, the packing of DNA by histones regulates the access of the cellular machinery to the underlying DNA. This is best exemplified by the fact that mutations that alter chromatin can result in transcription initiation from intragenic cryptic promoters in *S. cerevisiae* (Kaplan *et al*. 2003; Mason and Struhl 2003; Cheung *et al*. 2008). However, multiple cellular processes disrupt histone-DNA contacts, including elongation by DNA and RNA polymerases, and thus mechanisms must exist to reestablish chromatin structure during or following these events to maintain the fidelity of other cellular processes, such as transcription initiation.

Although elegant in vitro experiments suggest that RNA polymerase II can “step around” nucleosomes without displacing the histone octamer from DNA (Kulaeva *et al*. 2009; Chang *et al*. 2014), additional factors are required in vivo to maintain chromatin structure on transcribed genes. Key among these were identified through SuPpressor of Ty (SPT) screens in *S. cerevisiae*, which selected for mutants that restored the production of functional proteins from genes with δ insertions (Winston 1992). Of the proteins identified, Spt5, Spt6, and Spt16 co-localize with RNA polymerase II, consistent with roles in preventing histone displacement from transcribed genes (Mason and Struhl 2003; Mayer *et al*. 2010; Feng *et al*. 2016; Pathak *et al*. 2018; Martin *et al*. 2018). These proteins are well-conserved in eukaryotes, but the process of cryptic transcription in organisms with more complex genomes is only beginning to be explored. In a subsequent screen, the Winston Lab introduced a *HIS3* reporter downstream of a cryptic promoter in the *FLO8* gene and identified additional proteins involved in maintaining chromatin structure, including histone chaperones, chromatin remodelers, histone deacetylases, and components of the Mediator complex (Cheung *et al*. 2008). Cryptic transcription in cells lacking many of these factors depended on the activation of an upstream promoter, further supporting their role in preventing transcription-dependent histone displacement from the *FLO8* cryptic promoter.

Although transcription from an upstream promoter is required for the activation of cryptic promoters in some mutants, it is not strictly needed, including in *ASF1*, *SPT6*, *SPT16*, and *SPT21* mutants (Cheung *et al*. 2008), suggesting that a process other than transcription can destabilize nucleosomes. An obvious candidate is DNA replication, which is inherently disruptive to chromatin structure, and indeed, many replication-coupled factors, including Asf1, Rtt106, Spt6, and Spt16, are suppressors of cryptic transcription (Wittmeyer *et al*. 1999; Schlesinger and Formosa 2000; O’Donnell *et al*. 2004; Zhou and S-F Wang 2004; Groth *et al*. 2005, 2007; Franco *et al*. 2005; Schulz and Tyler 2006; VanDemark *et al*. 2006; Sanematsu *et al*. 2006; Han *et al*. 2007; Li *et al*. 2008a; Grigsby *et al*. 2009; Abe *et al*. 2011; Ishikawa *et al*. 2011; Zunder *et al*. 2012; Klimovskaia *et al*. 2014; Yang *et al*. 2016; Safaric *et al*. 2022; Nakagawa *et al*. 2022; Miller and Winston 2023). However, these proteins also have transcription-linked functions (Orphanides *et al*. 1998; Mason and Struhl 2003; Adkins *et al*. 2004; Biswas *et al*. 2005; Imbeault *et al*. 2008; Birch *et al*. 2009; Hsieh *et al*. 2013), and whether DNA replication is required for cryptic transcription following loss of these suppressor proteins has never been tested. Also noticeably absent from screens for suppressors of cryptic promoters are genes encoding the Chromatin Assembly Factor-1 (CAF-1) complex, a trimeric complex made up of Rlf2/Cac1, Cac2, and Msi1/Cac3. CAF-1 deposits histone H3-H4 tetramers onto nascent DNA during S phase, and loss of CAF-1 results in defects in chromatin assembly (Sauer *et al*. 2017; Panne *et al*. 2018). Thus, at this time, it is not clear whether nucleosome disruption due to DNA replication can lead to the activation of cryptic promoters.

In this work, we sought to test whether DNA replication can promote cryptic transcription in *S. cerevisiae*. Using a novel fluorescent reporter assay, we show that depletion of multiple factors, including Asf1, Rtt106, Spt6, and Spt16, results in activation of transcription from a cryptic promoter, but this is suppressed by the prior arrest of cells in the G1 phase of the cell cycle. This cell-cycle requirement for chromatin disruption, together with previous work linking Asf1, Rtt106, Spt6, and Spt16 to DNA replication, suggests that in the absence of these factors, passage through S phase can result in the activation of cryptic promoters. Additionally, we show that short-term depletion of the CAF-1 complex results in modest levels of cryptic transcription, but this effect is absent following long-term loss, suggesting the activation of a compensatory mechanism for chromatin assembly that maintains the fidelity of transcription initiation.

## Materials and Methods

### Yeast strains and growth conditions

All strains used in this study are listed in Table S1. The FRB tags were integrated at endogenous alleles by homologous recombination of PCR products. Strains with *spt16-197* or *spt6-1004* mutations or null alleles of *RLF2* or *RTT106* were created using CRISPR (https://benchling.com/pub/ellis-crispr-tools#reference). All strains were verified by PCR or sequencing of PCR products in the case of point mutants. Unless otherwise specified, yeast strains were grown at 30°C in YPD (yeast extract, peptone, 2% dextrose) and maintained at an OD_600_ of less than 1 for the entirety of the experiment. For G1 arrests, cells were incubated with 10 nM α factor for two hours. For anchor-away, rapamycin, dissolved to 8 μg/ml in DMSO, was added to a final concentration of 8 ng/ml, and incubation continued for two hours. Temperature-sensitive strains were incubated at the non-permissive temperature for two hours.

### Flow cytometry and statistical analyses

Cells were resuspended in cold phosphate-buffered saline, and forward scatter, side scatter, and fluorescence intensity (525/30 nm) of 10,000 cells were measured using a Guava® easyCyteTM Flow Cytometer. Subsequent data analyses were performed in R Studio. Median values of GFP signal from three biological replicates were calculated for each treatment and compared using a non-parametric one-tailed Mann-Whitney U test. The significance level was set to be 0.05, and any comparison that met this threshold (i.e., Ustat ≤ Ucrit) is described as significant in the text.

## Results

### Generation of a fluorescent reporter for cryptic transcription

To rapidly track cryptic transcription in *S. cerevisiae*, we followed a similar approach as Cheung et al., except inserted a fluorescent sfGFP reporter downstream of a cryptic promoter in the *FLO8* gene (Figure 1A). The insertion site placed the sfGFP ORF in a +1-frameshift relative to the *FLO8* start codon such that only transcripts originating from the cryptic promoter, and not the *FLO8* promoter, were translated into a fluorescent gene product. To test our reporter system, we chose a well-characterized mutation in *SPT16*, which encodes a component of the FACT complex. Previous work has shown that in this mutant, *spt16-167*, Spt16 is rapidly degraded at the non-permissive temperature (37°C), resulting in genome-wide loss of nucleosome occupancy and positioning (Dasgupta *et al*. 2004; van Bakel *et al*. 2013; Feng *et al*. 2016). We therefore incubated cells at the non-permissive temperature and used flow cytometry to track the appearance of GFP signal at varying intervals following the temperature shift. While wild-type cells showed a minimal level of fluorescence (Figure 1B), *spt16-197* cells exhibited a significant time-dependent increase in GFP signal (Figure 1C), suggesting the reporter can be used to detect cryptic transcription in single cells.

**Figure 1.**
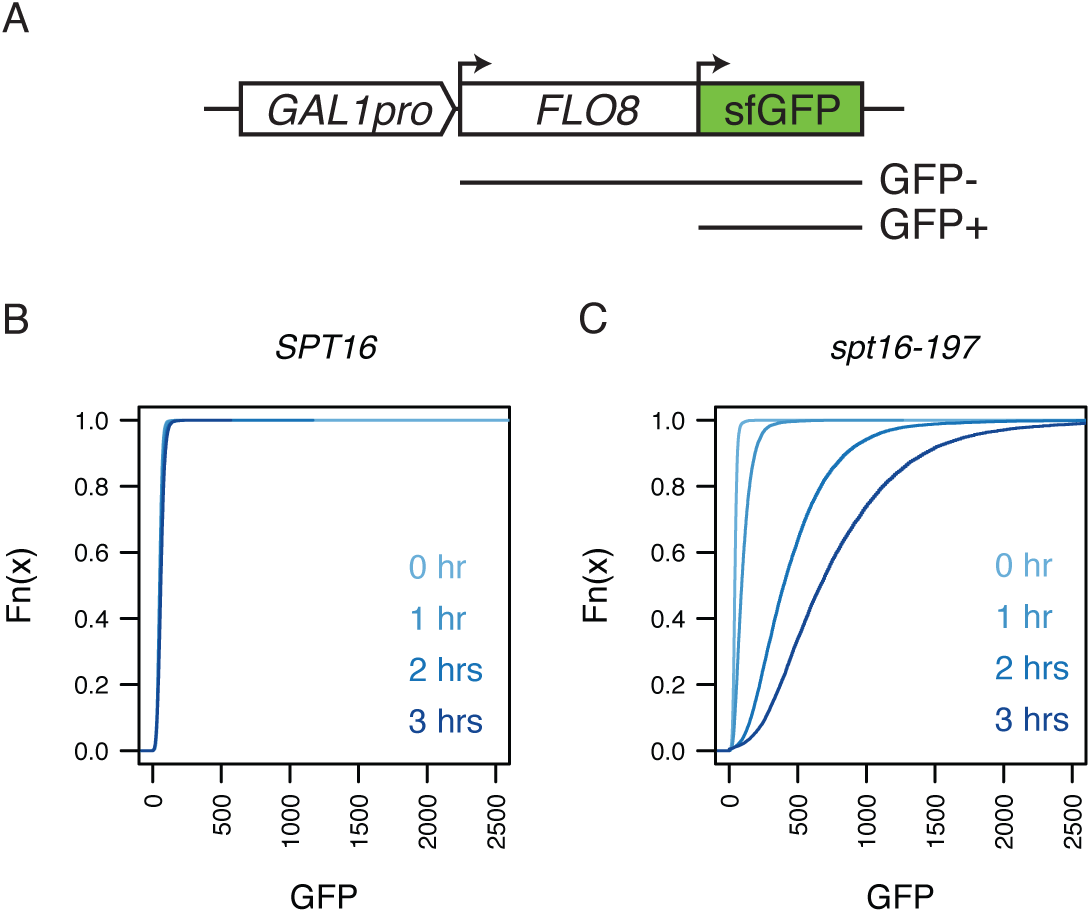
Development of a fluorescent reporter for cryptic transcription. A) Schematic of the *FLO8:sfGFP* reporter. The sfGFP open reading frame was inserted downstream of the cryptic promoter with a +1-frameshift relative to the *FLO8* start codon. The length of the expected full-length and cryptic transcripts, as well as the GFP phenotypes, are indicated below. The *GAL1* promoter regulated *FLO8* expression for all experiments except those shown in Figures 1B and C, in which the endogenous *FLO8* promoter was used. B and C) Empirical cumulative distribution function plots of GFP signal measured by flow cytometry of 10,000 wild type (*SPT16*) and *spt16-197* cells grown at the non-permissive temperature (37°C) for the indicated times.

The *spt16-197* mutant is reported to have a cell-cycle defect at the non-permissive temperature. We reanalyzed the data using a subpopulation of similarly sized cells from each sample to rule out the possibility that cell size differences between the wild-type and *spt16-197* populations were driving the altered GFP signal. The results were unchanged (Figures S1A and B). To ensure that the GFP signal was not due to the increased time spent in G1, we treated wild-type cells carrying the sfGFP reporter with α factor to induce a similar arrest. Wild-type-arrested cells exhibited an equivalent level of background fluorescence signal as asynchronous cells (Figure S1C). We, therefore, concluded that the effect of cell-cycle arrest on the GFP signal could be taken out of consideration for further experiments, and our reporter could be used to detect cryptic transcription.

### A fluorescent reporter of cryptic transcription can differentiate between chromatin disruption due to transcription and DNA replication

Transcription and DNA replication are two cellular processes that destabilize chromatin. Using a regulatable *GAL1pr-FLO8-HIS3* reporter, Cheung *et al*. showed that, in a subset of mutants, transcription from an upstream reporter was required for activation of the *FLO8* cryptic promoter, but the growth requirement of the *HIS3* reporter prevented them from testing whether DNA replication was required (Cheung *et al*. 2008). Because our reporter assay is not dependent on cell growth to detect cryptic transcription, we were able to directly test whether passage through S phase promotes activation of cryptic promoters following depletion of chromatin stabilizing factors. We began by depleting Spt21 from yeast nuclei using anchor-away (Haruki *et al*. 2008). Spt21 is required to induce the histone genes in S phase, and deletion of *SPT21* results in transcription initiation from cryptic promoters (Dollard *et al*. 1994). The likely explanation for this is that in the absence of Spt21, cells cannot synthesize sufficient histones in S phase to package nascent chromatin properly. Assays for cryptic transcription using our fluorescent reporter were performed in biological triplicate, and the median GFP signals are shown in Figure 2A, with the cumulative distribution plots shown in Figure S2. To prevent upstream-originating transcription through the cryptic promoter, we replaced the *FLO8* promoter with a *GAL1* promoter and grew the cells on dextrose. Depletion of Spt21 resulted in increased GFP signal in greater than 80% of cells, indicating a highly penetrant cryptic transcription phenotype (Figure S2). However, arresting cells in G1 with α factor before depleting Spt21 rescued this phenotype, and thus passage through the cell cycle was required for cryptic transcription following depletion of this protein. Similar results were observed following the anchor-away of Spt10, a second factor necessary for full histone gene induction in S phase (Figures 2B and S3) (Dollard *et al*. 1994). In contrast, depletion of Spt2 in cells grown in dextrose did not result in GFP production in either synchronous or asynchronous cells (Figure 2C and S4), however, a GFP signal was observed when cells were grown on galactose, and this was not reduced by G1 arrest (Figure 2D and S5), consistent with a role for Spt2 in stabilizing chromatin disrupted by RNA polymerase (Chen *et al*. 2015). These results collectively suggest that defects in chromatin assembly during S phase can activate cryptic promoters. Further, our reporter can be used to determine whether chromatin disruption is due to the activity of DNA or RNA polymerases.

**Figure 2.**
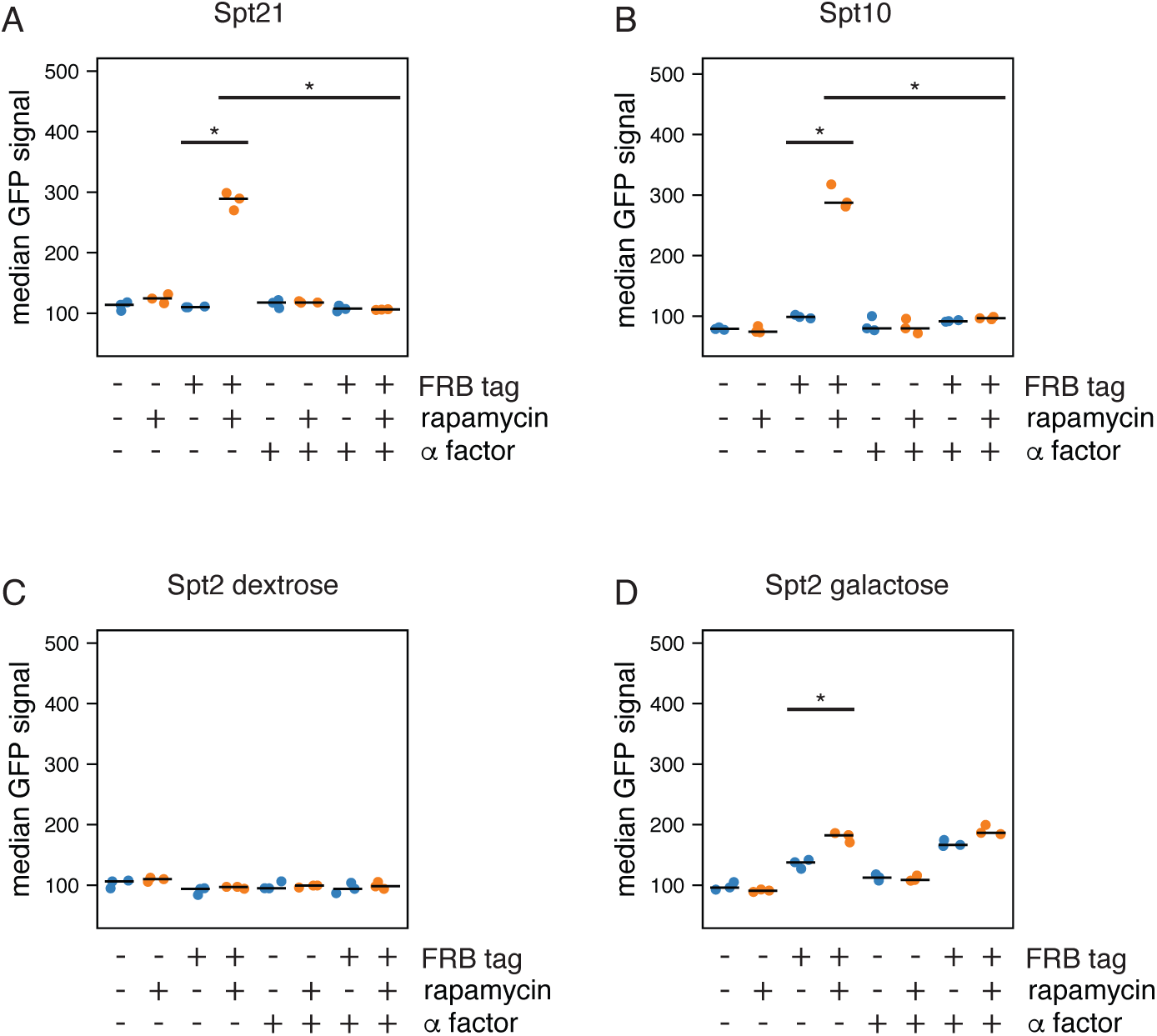
Cell cycle arrest suppresses cryptic transcription caused by Spt10 or Spt21 depletion. A-D) The indicated proteins were depleted using anchor-away by the addition of a C-terminal FRB tag and treatment with rapamycin (orange), and transcription from a *FLO8*-sfGFP cryptic promoter was assessed by flow cytometry of 10,000 cells. Cultures lacking rapamycin were treated with DMSO as vehicle control (blue). Cells were either asynchronous or arrested in G1 with α factor prior to rapamycin addition. Assays were performed on three independent cultures and the median GFP signal of each culture was plotted. “*” indicates a statistically significant difference using a Mann-Whitney U test (alpha = 0.5, one-tailed).

### The CAF-1 complex suppresses transcription initiation from a cryptic promoter

As suggested above, proper chromatin assembly on newly replicated DNA is vital to suppress transcription initiation from aberrant sites. The CAF-1 complex plays a well-characterized role in depositing histones H3 and H4 on nascent DNA (Sauer *et al*. 2017; Panne *et al*. 2018), but surprisingly, genes encoding this complex have not been identified in screens for suppressors of cryptic transcription. Figure 3A, which used the *GAL1pr-FLO8-HIS3* reporter to examine the cryptic transcription phenotype following loss of *RLF2/CAC1,* confirmed this result. To determine whether we could detect a cryptic transcription phenotype due to the loss of CAF-1 using our reporter assay, we depleted Rlf2/Cac1 using anchor away. Figures 3B and S6 show that loss of Rlf2/Cac1 from the nucleus resulted in a statistically significant increase in GFP signal, which was suppressed by arresting cells in G1, consistent with a documented function for CAF-1 in assembling chromatin on nascent DNA. However, the effect was subtle (note that the Y-axis range is 50% of that in Figure 2). One explanation for the modest impact was inefficient anchor-away, so we repeated the analysis using a null allele of *RLF2/CAC1*. Surprisingly, a *rlf2/cac1*11 mutant showed a reduced effect (compare Figures 3C and D). These results indicate that while short-term loss of CAF-1 can destabilize chromatin on nascent DNA, a mechanism exists to counteract the effects of long-term depletion. Additionally, these results explain why genes encoding subunits of CAF-1 were not identified in screens for suppressors of cryptic transcription.

**Figure 3.**
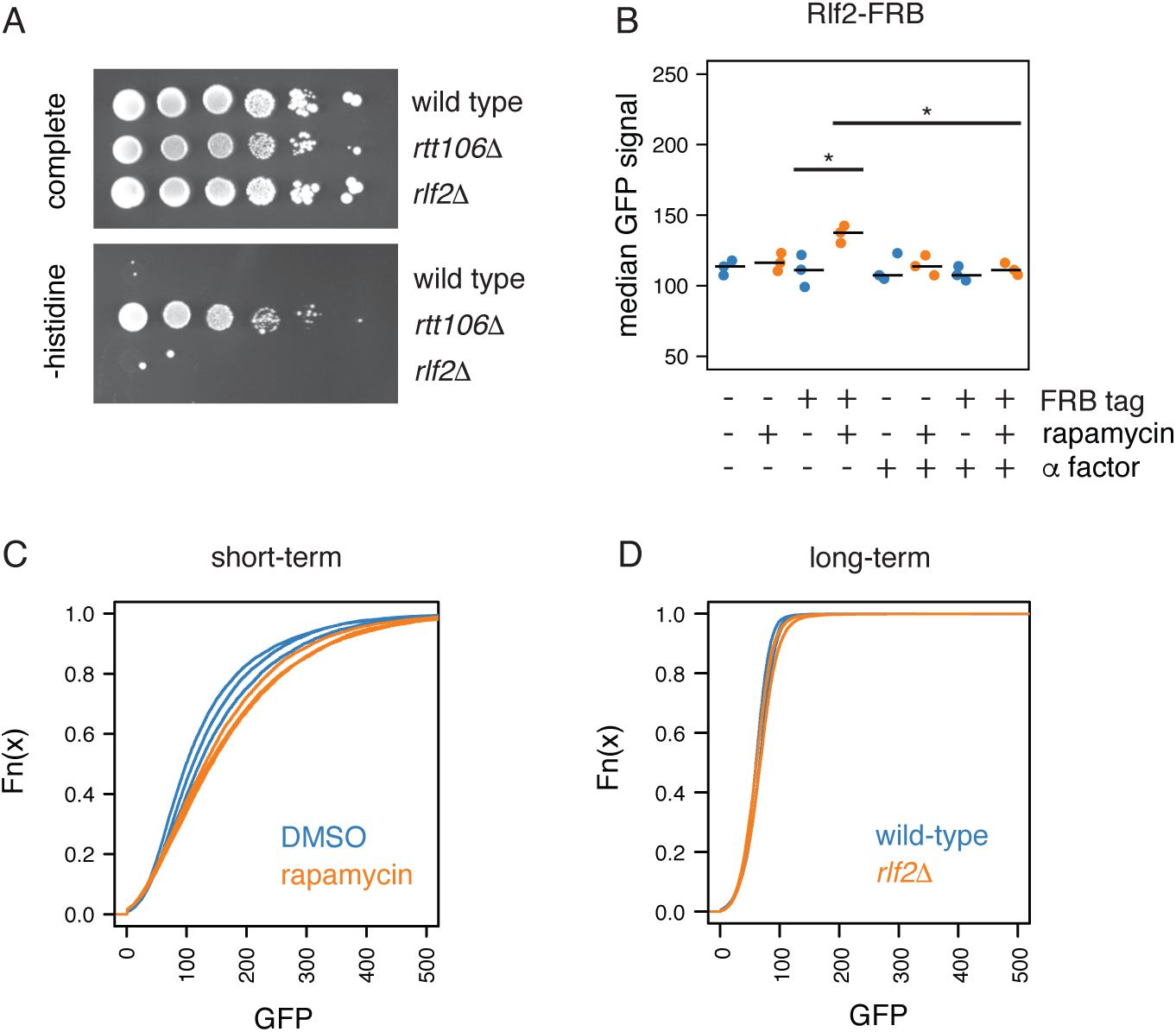
Short-term depletion of CAF-1 results in cryptic transcription. A) Ten-fold serial dilutions of the indicated *FLO8::HIS3* strains were plated on synthetic complete or histidine drop-out media and incubated at 30°C for three days. B) Rlf2-FRB was depleted using anchor-away (orange) from a *FLO8:sfGFP* strain, and the GFP levels were assessed by flow cytometry of 10,000 cells. Cultures lacking rapamycin were treated with DMSO as a vehicle control (blue). Cells were either asynchronous or arrested in G1 with α factor prior to rapamycin addition. Assays were performed on three independent cultures and the median GFP of each culture was plotted. “*” indicates a statistically significant difference using a Mann-Whitney U test (alpha = 0.5, one-tailed). C and D) GFP levels from asynchronous cells following short-term [anchor away (C)] or long-term [gene deletion (D)] of Rlf2.

### Confirming roles for transcription-linked chromatin-stabilizing factors in DNA replication

Multiple factors play dual functions in stabilizing chromatin structure on both transcribed and newly replicated DNA, including Asf1, Rtt106, Spt6, and FACT (Wittmeyer *et al*. 1999; Schlesinger and Formosa 2000; O’Donnell *et al*. 2004; Zhou and S-F Wang 2004; Groth *et al*. 2005, 2007; Franco *et al*. 2005; Schulz and Tyler 2006; VanDemark *et al*. 2006; Sanematsu *et al*. 2006; Han *et al*. 2007; Li *et al*. 2008a; Grigsby *et al*. 2009; Abe *et al*. 2011; Ishikawa *et al*. 2011; Zunder *et al*. 2012; Klimovskaia *et al*. 2014; Yang *et al*. 2016; Safaric *et al*. 2022; Nakagawa *et al*. 2022; Miller and Winston 2023), and we used our reporter assay to test whether some of the chromatin disruption that occurs in the absence of these factors is due to DNA replication. Consistent with previous work, depletion of Asf1 in cells grown in dextrose resulted in cryptic transcription that was entirely suppressed by G1 arrest, suggesting that at a non-transcribed locus, chromatin disruption in the absence of Asf1 is due to DNA replication (Figures 4A and S7). Anchor away of Rtt106 did not statistically affect the median GFP signal (Figure 4B), but interestingly, a cumulative distribution plot of the same data revealed a shift in the upper 50% of the distribution (Figures 4C and S8), and this was even more apparent in a *rtt106*11 strain (Figure 4D). While some of the distribution of the “background” GFP signal in wild-type *GAL1pr-FLO8-sfGFP* cells was likely due to inherent variation in the data, some may reflect the fact that a subpopulation of wild-type cells exhibits low levels of cryptic transcription. If this is the case, then the shift of the upper portion of the distribution curve may suggest that loss of Rtt106 exacerbates chromatin disruption in cells already undergoing low levels of cryptic transcription.

**Figure 4.**
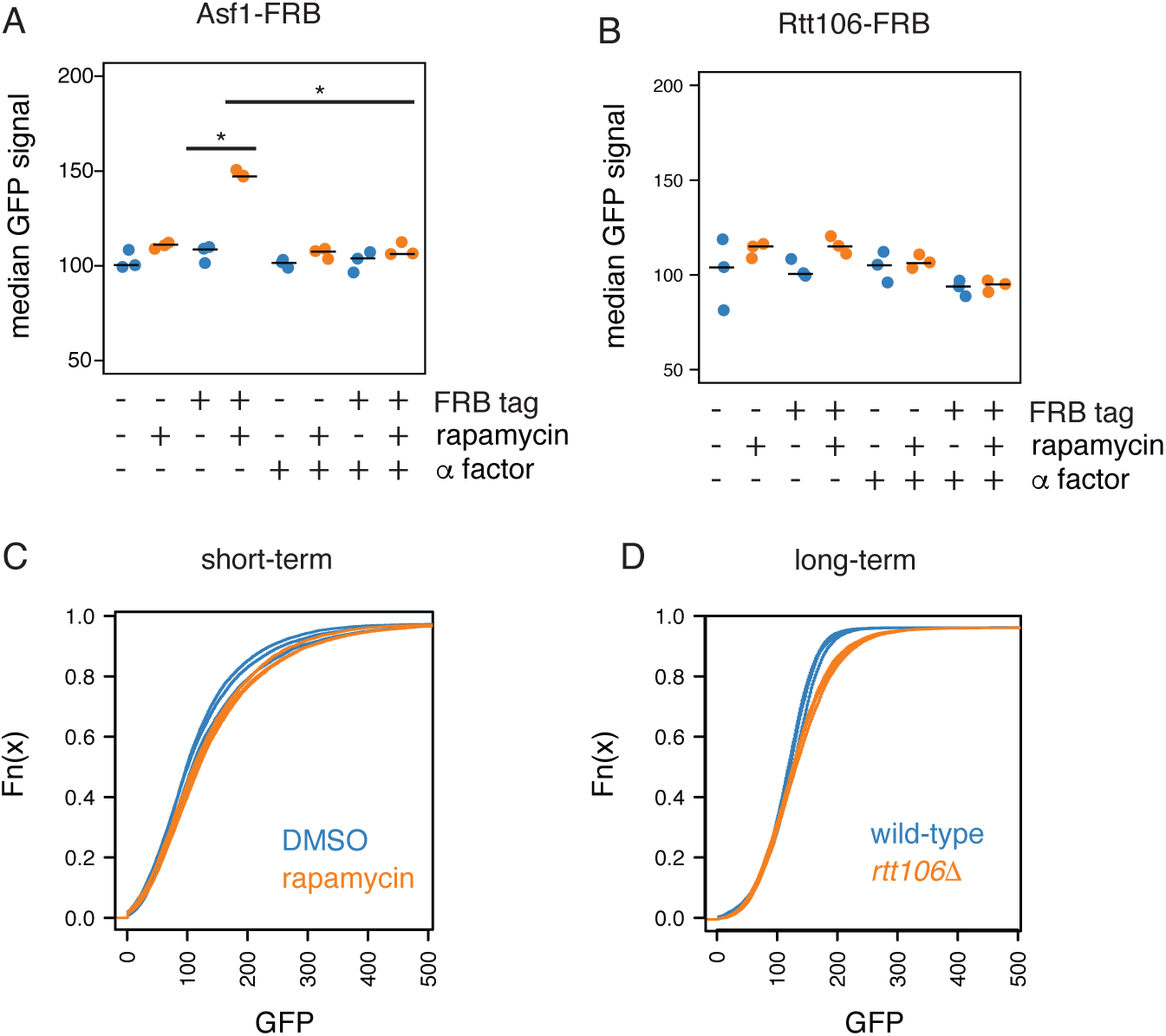
Cell cycle arrest suppresses cryptic transcription caused by Asf1 depletion. A and B) Asf1 (A) and Rtt106 (B) were depleted using anchor-away from a *FLO8:sfGFP* strain, and GFP levels assessed by flow cytometry of 10,000 cells. Cells were either asychronous, or arrested in G1 with α factor prior to rapamycin addition. Cultures lacking rapamycin were treated with DMSO as a vehicle control. Assays were performed on three independent cultures and the median GFP of each culture plotted. “*” indicates a statistically significant difference using a Mann-Whitney U test (alpha = 0.5, one-tailed). C and D) GFP levels from asychronous cells following short-term [anchor away (C)] or long-term [gene deletion (D)] of Rtt106.

Numerous studies have demonstrated a role for the FACT complex in stabilizing chromatin on nascent DNA. To further support this, we repeated our analysis with the *spt16-197* temperature-sensitive mutant in cells grown in dextrose, with and without treatment with α factor. Figures 5A and S9 show that a G1 arrest before the shift to the non-permissive temperature partially suppressed the cryptic transcription phenotype, suggesting that some of the chromatin disruption in the absence of FACT was due to DNA replication. Similar results were observed in a temperature-sensitive mutant of *SPT6* (Figures 5B and S10), which encodes a histone chaperone recently implicated in DNA replication (Miller and Winston 2023). Because both *SPT6* and *SPT16* mutants were grown on dextrose, which represses transcription from the upstream *GAL1* promoter, the incomplete rescue implies that depletion of FACT or Spt6 results in chromatin disruption in the absence of DNA replication and transcription. However, loss of Spt6 or Spt16 has been shown to result in genome-wide aberrant transcription initiation (Cheung *et al*. 2008; Feng *et al*. 2016; Doris *et al*. 2018), and thus cryptic transcription from either upstream or downstream sites could disrupt chromatin over the *FLO8* cryptic promoter. Regardless of the cause, the decreased cryptic transcription in non-cycling cells supports the importance of Spt6 and Spt16 in stabilizing chromatin on newly replicated DNA.

**Figure 5.**
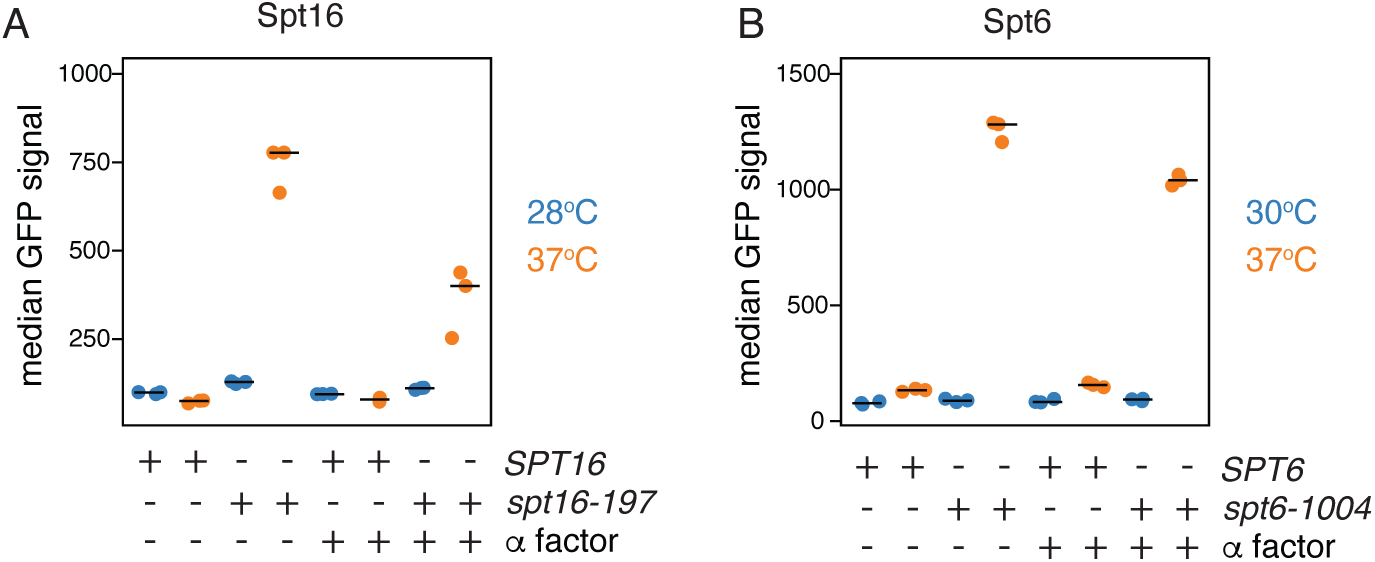
Cell cycle arrest partially rescues cryptic transcription defects in *SPT16* and *SPT6* mutants. A and B) The indicated mutants were grown at permissive (blue) or non-permissive (orange) temperatures and transcription from a *FLO8*-sfGFP cryptic promoter assessed by flow cytometry of 10,000 cells. Cells were either asynchronous, or arrested in G1 with α factor prior to a shift to the non-permissive temperature. Assays were performed on three independent cultures and the median GFP of each culture was plotted. “*” indicates a statistically significant difference using a Mann-Whitney U test (alpha = 0.5, one-tailed).

## Discussion

In this work, we developed a novel assay for cryptic transcription with additional benefits compared to other approaches. First, because it is not a growth-based assay, it allowed us to reveal a requirement for cell-cycle passage for cryptic transcription following depletion of Asf1, Spt6, Spt10, Spt21, or the CAF-1 and FACT complexes. This feature also facilitated the assessment of cryptic transcription following the short-term absence of factors. This was critical in the case of Rlf2/Cac1, the loss of which results in cryptic transcription following short-term, but not long-term, depletion. Second, because the assay employs single cells, it can be used to determine whether all cells or just a subpopulation are exhibiting cryptic transcription. Using this feature, we revealed that depletion of multiple proteins required for proper chromatin assembly, such as Asf1, Spt6, Spt10, Spt16, and Spt21, resulted in cryptic transcription in over 80% of cells, suggesting it is not a rare event in these mutants. A weakness of the assay, however, is that it requires a well-characterized cryptic transcription start site to drive the expression of the reporter in a different reading frame from the protein-coding gene it is embedded in. Fortunately, the data from a recent genome-wide survey identified over 6000 sense-strand cryptic transcription start sites in a *spt6-1004* mutant (Doris *et al*. 2018). Thus, many additional sites can be investigated.

Using our reporter, we generated data suggesting that defects in chromatin assembly during S phase can activate cryptic promoters. Several studies have shown that on newly replicated DNA, nucleosome spacing across gene bodies is altered, and a “maturation” step is required before regular phasing is observed (Vasseur *et al*. 2016; Fennessy and Owen-Hughes 2016; Gutiérrez *et al*. 2019). Further, the loss of *CAC1* or *ASF1* delays this step (Fennessy and Owen-Hughes 2016). Because cryptic promoters share similar sequence features as canonical promoters but are depleted in binding sites for factors that typically exclude nucleosomes (Doris *et al*. 2018), a delay in chromatin maturation could provide an opportunity for the transcriptional machinery to access these sites. Surprisingly, depletion of Asf1, Spt10, or Spt21 led to a GFP signal in over 80% of the cells in a population, and this was entirely suppressed by arresting cells in G1. Considering this ubiquitous loss of transcription fidelity during chromatin replication, it is surprising that *asf1*11, *spt10*11, and *spt21*11 mutants are viable. However, although most cells showed some GFP signal, the amount was reduced compared to the loss of essential factors, such as Spt6 and Spt16. A likely explanation is that decreases in histone levels following depletion of Asf1, Spt10, or Spt21 delay but does not impede chromatin maturation, resulting in only transient activation of the cryptic reporter.

Despite the well-characterized roles of CAF-1 and Rtt106 in assembling chromatin on nascent DNA, depletion of these factors resulted in relatively modest levels of cryptic transcription compared to loss of others such as Asf1, Spt6, or Spt16. A possible explanation for this is that CAF-1 and Rtt106 play redundant roles in S phase, and there is support for this in the literature. CAF-1 and Rtt106 have both been reported to accept histones H3 and H4 from Asf1 for deposition on DNA (Li *et al*. 2008b; Fazly *et al*. 2012; Han *et al*. 2013; Mattiroli *et al*. 2017) and multiple high-throughput studies have reported synthetic growth defects in strains lacking both Rtt106 and CAF-1. Such a redundancy could explain another intriguing observation made in this study: the transient effect of CAF-1 depletion on cryptic transcription. Because *RTT106* expression peaks in S phase (Spellman *et al*. 1998), a prolonged S phase due to the absence of CAF-1 could result in an increased window of *RTT106* expression, masking the long-term depletion of CAF-1. Thus, it would be interesting to test whether the combined anchor-away of these factors enhances the cryptic transcription phenotype. An alternate explanation for the limited impact of CAF-1 and Rtt106 depletion on cryptic transcription is that the *FLO8* cryptic promoter is less sensitive to the loss of these factors than others. Indeed, recent work has shown that the delay to chromatin maturation in a *CAC1* null mutant is not uniform for all nucleosomes, with some nucleosomes maturing with wild-type kinetics (Chen *et al*.). Since the timing of nucleosome maturation impacts the window of opportunity for the transcriptional machinery to access promoters, this variable could dictate whether a cryptic promoter is sensitive to CAF-1 loss. In contrast, depletion of Spt16, which has been shown to more negatively impact bulk histones levels than the absence of CAF-1 (van Bakel *et al*. 2013), could impact more nucleosomes, activating more cryptic promoters.

While our results suggest that chromatin disruption due to DNA replication can activate cryptic promoters, we also provide evidence for cell cycle-independent chromatin destabilization following the loss of Spt2, Spt6, and Spt16. In the case of Spt2, cryptic transcription depended on activating an upstream promoter suggesting this protein functions to stabilize chromatin on transcribed genes. Structural studies have shown that a single Spt2 molecule interacts with a histone H3-H4 tetramer, an interaction that on its own cannot shield all of the histone charges (Chen *et al*. 2015). Thus, we propose that the remaining charges are likely to be neutralized by DNA, and Spt2 plays a role akin to FACT by binding exposed histones on RNA polymerase-destabilized nucleosomes, preventing displacement by other chaperones (Martin *et al*. 2018; Liu *et al*. 2020; Zhou *et al*. 2020; Formosa and Winston 2020). In contrast to Spt2, depletion of Spt6 and Spt16 resulted in high levels of cryptic transcription in the absence of upstream promoter activation, and this was only modestly suppressed by G1 arrest suggesting the majority of chromatin destabilization is not due to DNA replication or transcription. However, loss of Spt6 and Spt16 results in transcription initiation from thousands of cryptic promoters (Cheung *et al*. 2008; Feng *et al*. 2016; Doris *et al*. 2018), and RNA polymerase originating from one of these promoters could destabilize chromatin over the *FLO8* cryptic promoter. Alternatively, a transcription and replication-independent pathway for nucleosome displacement over cryptic promoters in FACT and Spt6 mutants could exist. An interesting feature of canonical promoters is the transcription-independent turnover of histones in the flanking nucleosomes (Dion *et al*. 2007; Rufiange *et al*. 2007). Considering cryptic promoters share many of the features of canonical promoters, it would be interesting to test whether the nucleosomes flanking these promoters are more prone to turnover when FACT and Spt6 are absent. This and other insights revealed in this study will be the subject of future investigation.

## Data Availability

Strains are available upon request. The authors affirm that all data necessary to confirm the article’s conclusions are present in the article, figures, and tables.

## Acknowledgments

We are grateful to Fred Winston for providing strain FY2173 and the Sadowski and Ciernia Laboratories at UBC for technical support and advice for data analyses.

---

**Table S1.**
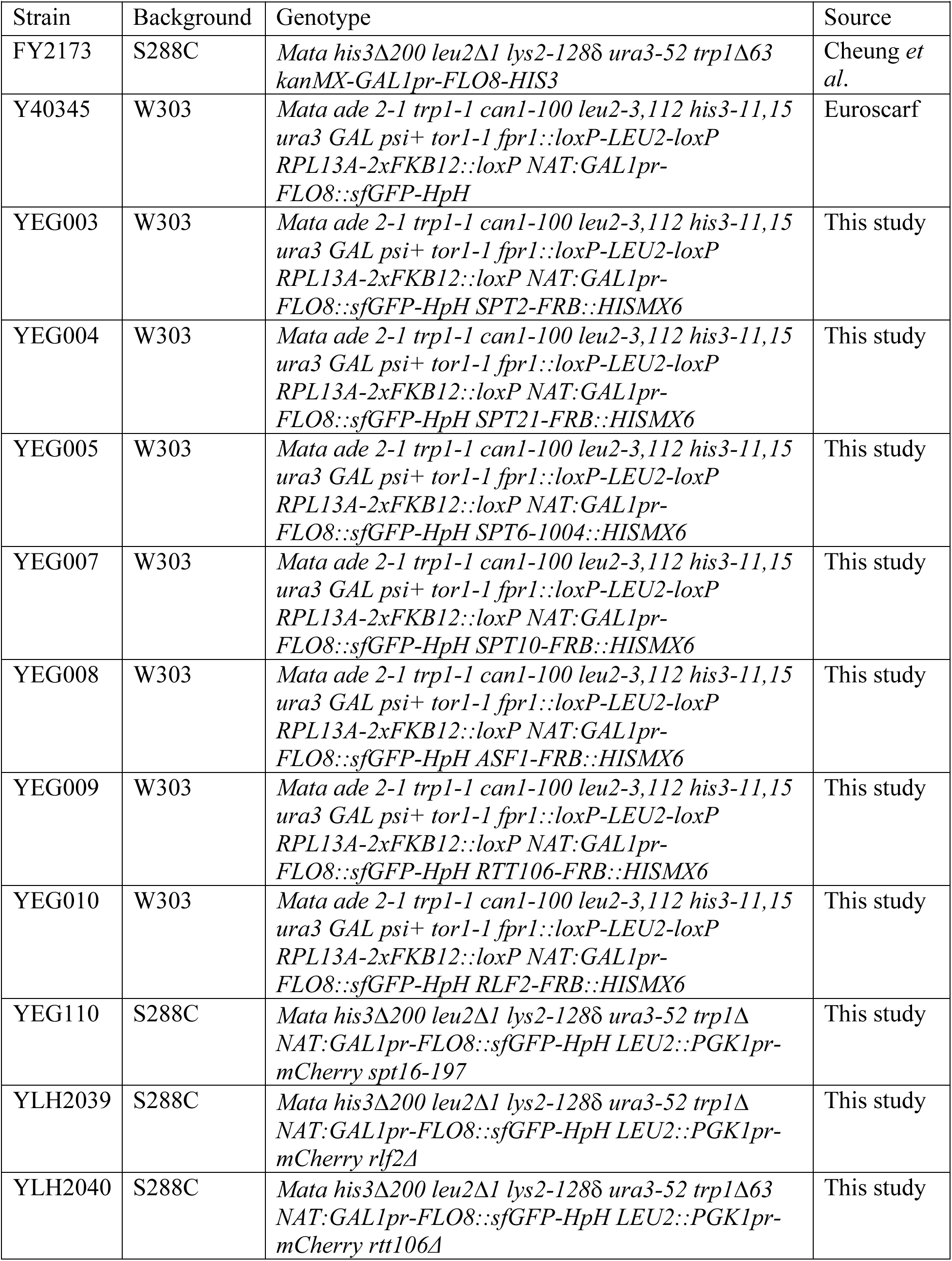
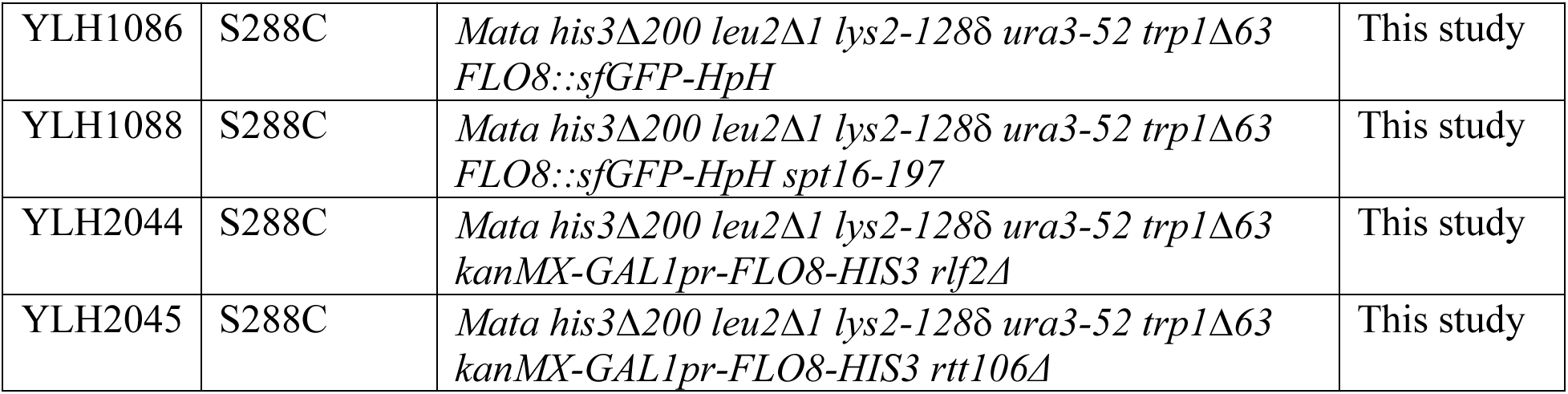
Strains used in this study

**Figure S1.**
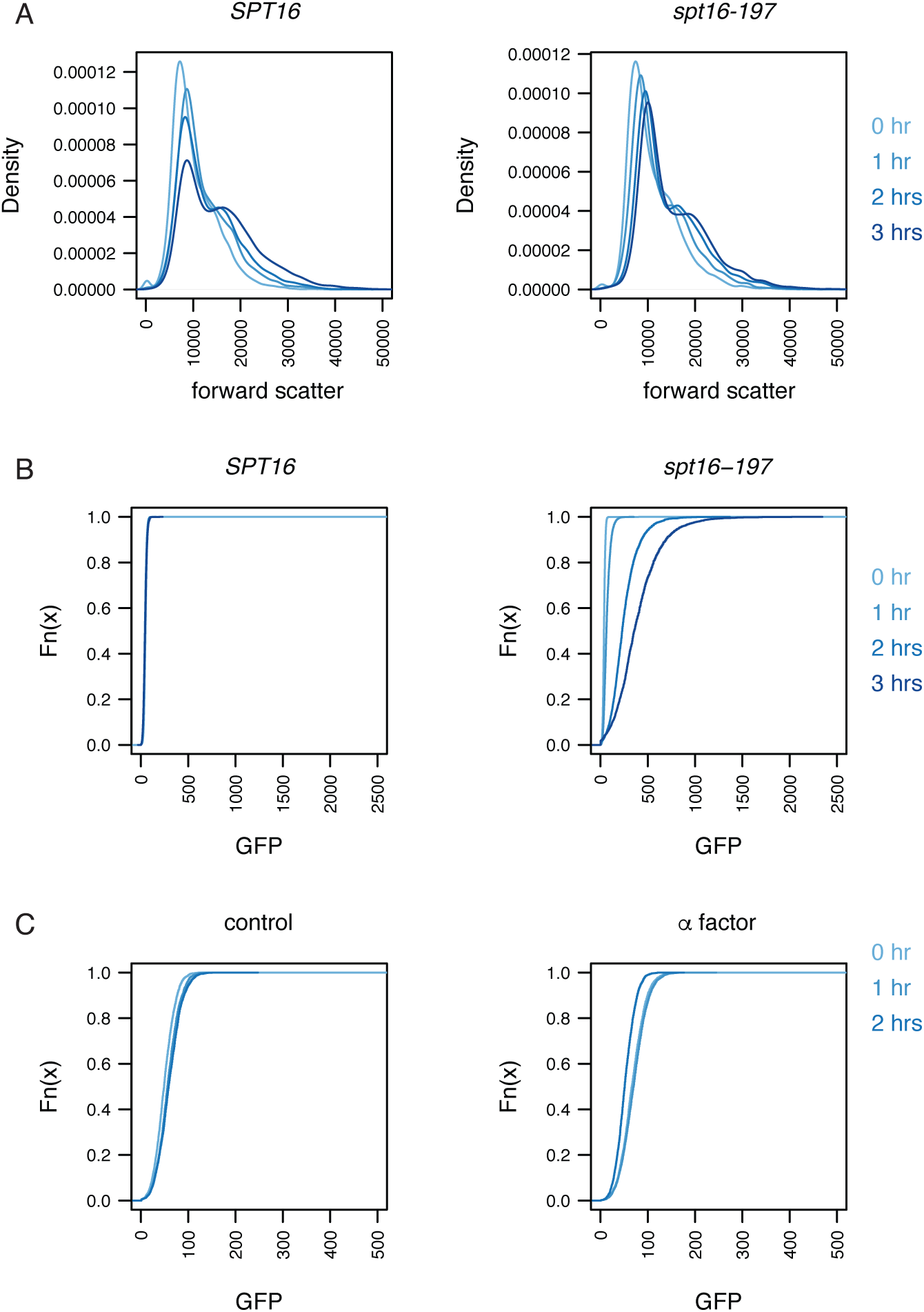
Cell cycle arrest does not impact the differential signals from a fluorescent reporter for cryptic transcription. A) Density plot of forward scatter of 10,000 wild-type (*SPT16*) and *spt16-197* cells grown at a non-permissive temperature (37°C) for the indicated times. B) Empirical cumulative distribution function plots of GFP signals from wild-type (*SPT16*) and *spt16-197* cells grown at a non-permissive temperature (37°C) for the indicated times and with a forward scatter of less than 10,000. C) Empirical cumulative distribution function plots of GFP signal measured by flow cytometry of 10,000 wild-type cells with and without α factor treatment for the indicated times.

**Figure S2.**
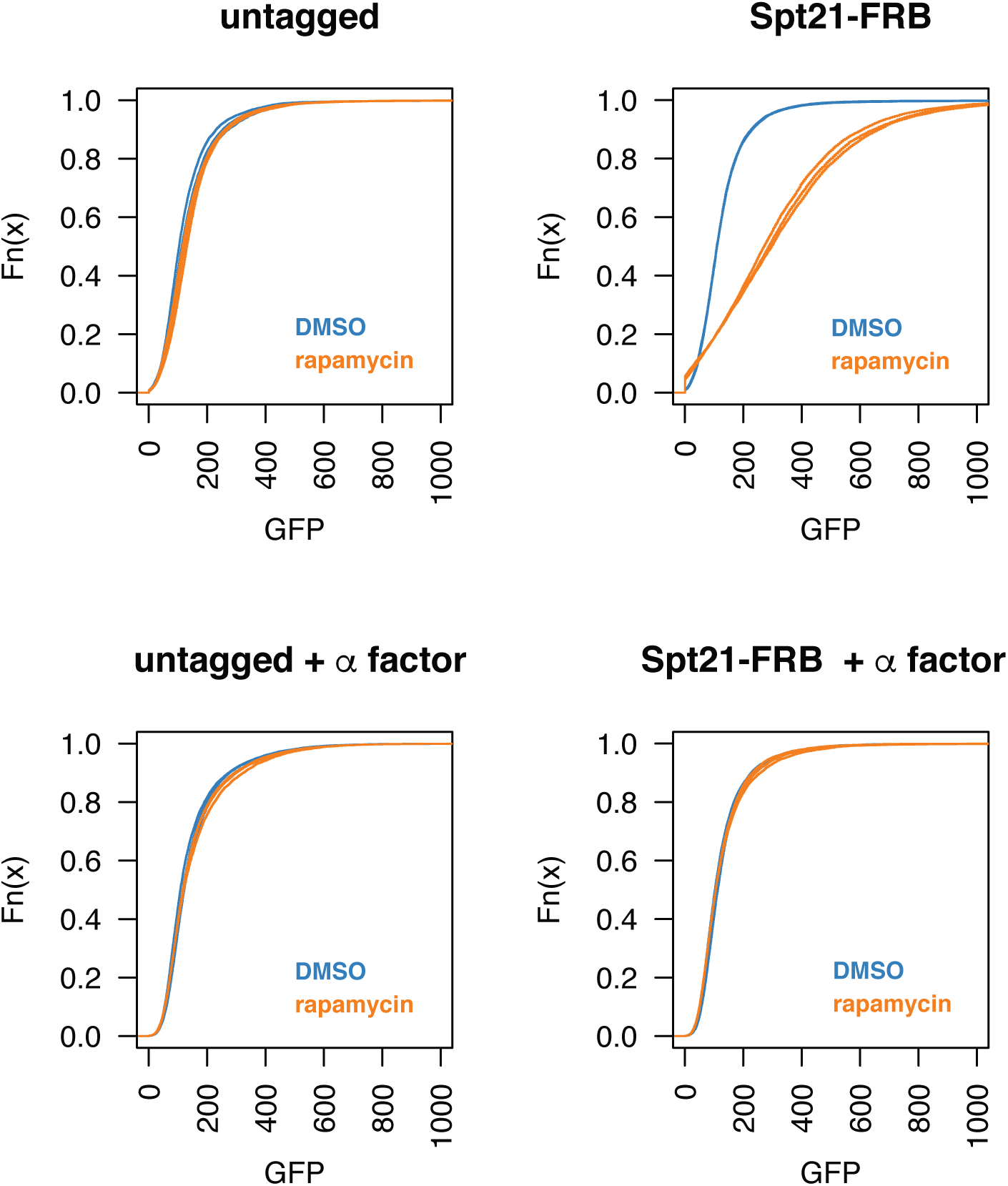
Cell cycle arrest rescues cryptic transcription defects due to depletion of Spt21. Spt21 was depleted using anchor-away by the addition of a C-terminal FRB tag and treatment with rapamycin (orange), and transcription from a *FLO8-sfGFP* cryptic promoter was assessed by flow cytometry of 10,000 cells. Cultures lacking rapamycin were treated with DMSO as vehicle control (blue). Cells were grown in dextrose and were either asynchronous or arrested in G1 with α factor prior to rapamycin/DMSO addition. Assays were performed on three independent cultures and empirical cumulative distribution function plots of each are presented.

**Figure S3.**
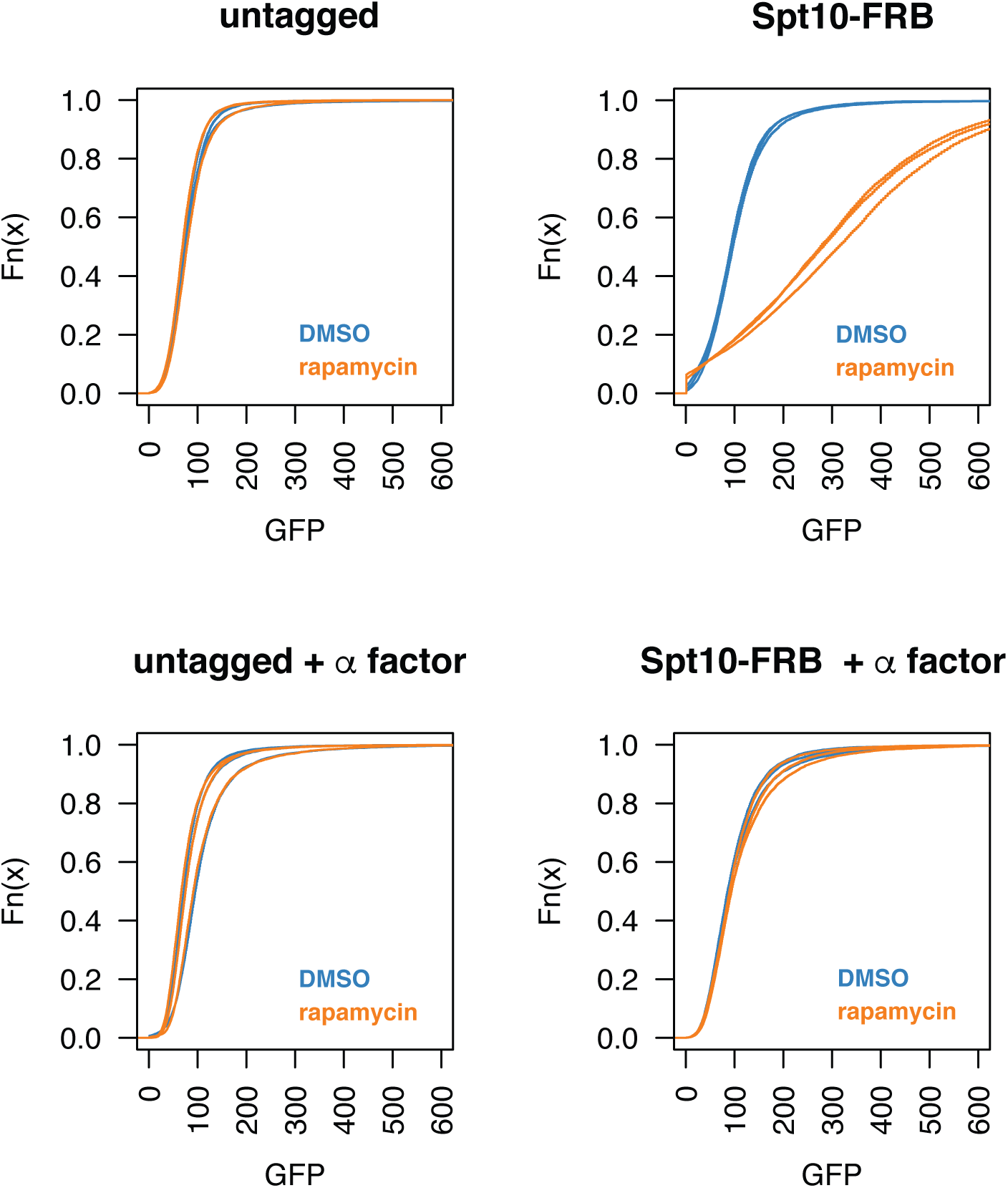
Cell cycle arrest rescues cryptic transcription defects due to depletion of Spt10. Spt10 was depleted using anchor-away by the addition of a C-terminal FRB tag and treatment with rapamycin (orange), and transcription from a *FLO8-sfGFP* cryptic promoter was assessed by flow cytometry of 10,000 cells. Cultures lacking rapamycin were treated with DMSO as vehicle control (blue). Cells were grown in dextrose and were either asynchronous or arrested in G1 with α factor prior to rapamycin/DMSO addition. Assays were performed on three independent cultures and empirical cumulative distribution function plots of each are presented.

**Figure S4.**
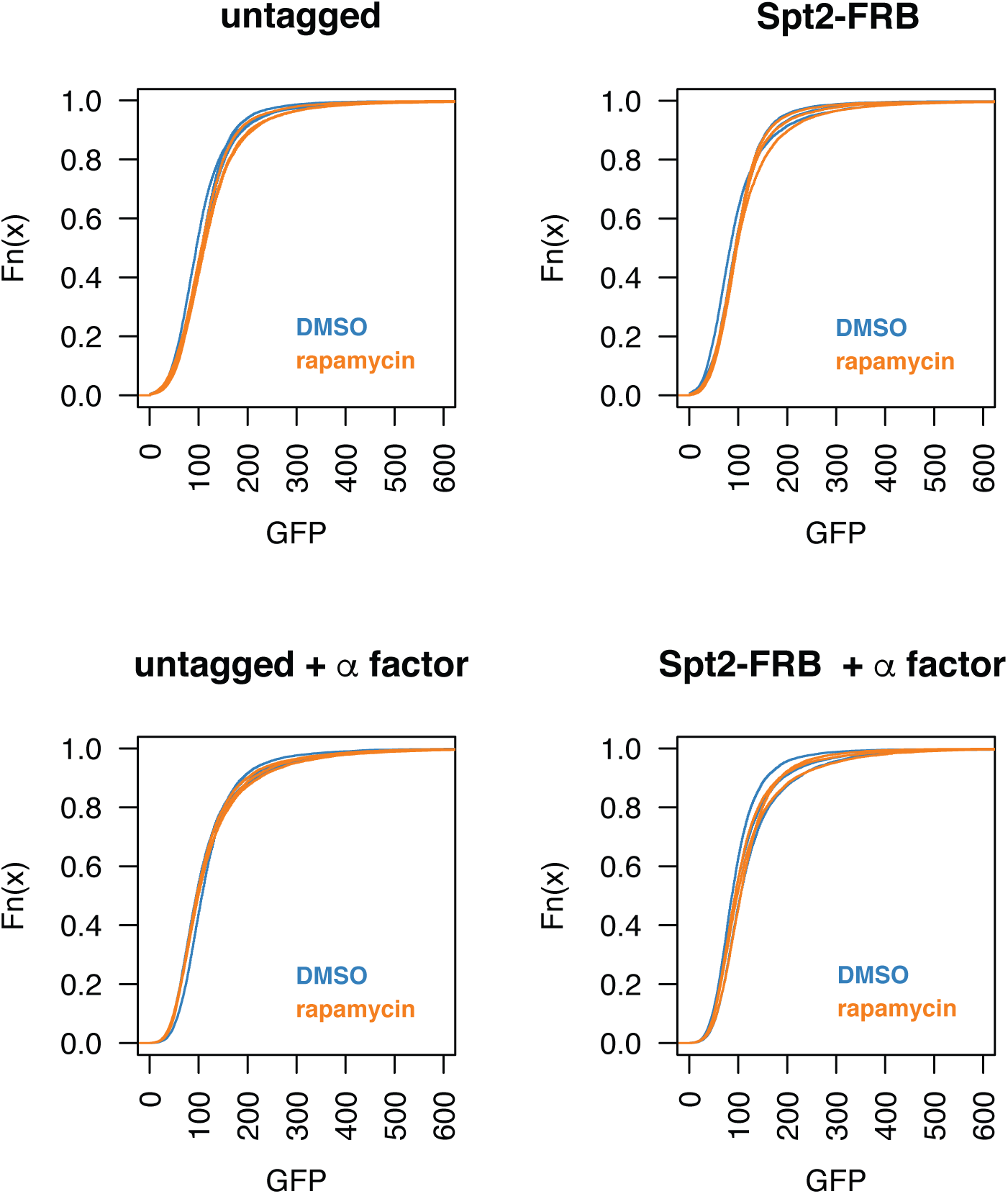
Depletion of Spt2 does not activate a cryptic promoter located within a non-transcribed gene. Spt2 was depleted using anchor-away by the addition of a C-terminal FRB tag and treatment with rapamycin (orange), and transcription from a *FLO8-sfGFP* cryptic promoter was assessed by flow cytometry of 10,000 cells. Cultures lacking rapamycin were treated with DMSO as vehicle control (blue). Cells were grown in dextrose and were either asynchronous or arrested in G1 with α factor prior to rapamycin/DMSO addition. Assays were performed on three independent cultures and empirical cumulative distribution function plots of each are presented.

**Figure S5.**
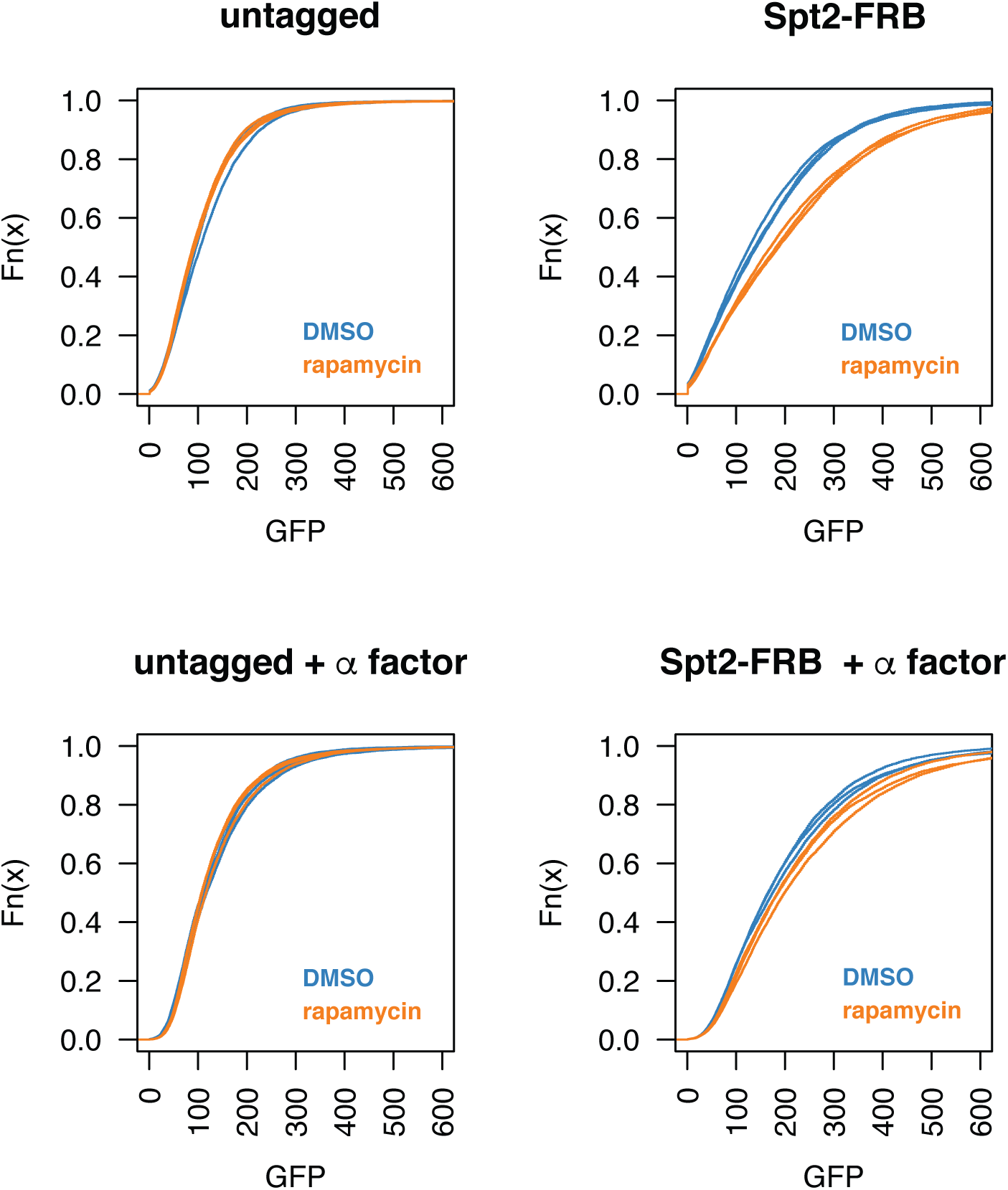
Transcription promotes activation of a cryptic promoter following depletion of Spt2. Spt2 was depleted using anchor-away by the addition of a C-terminal FRB tag and treatment with rapamycin (orange), and transcription from a *FLO8-sfGFP* cryptic promoter was assessed by flow cytometry of 10,000 cells. Cultures lacking rapamycin were treated with DMSO as a vehicle control (blue). Cells were grown in galactose and were either asynchronous or arrested in G1 with α factor prior to rapamycin/DMSO addition. Assays were performed on three independent cultures and empirical cumulative distribution function plots of each are presented.

**Figure S6.**
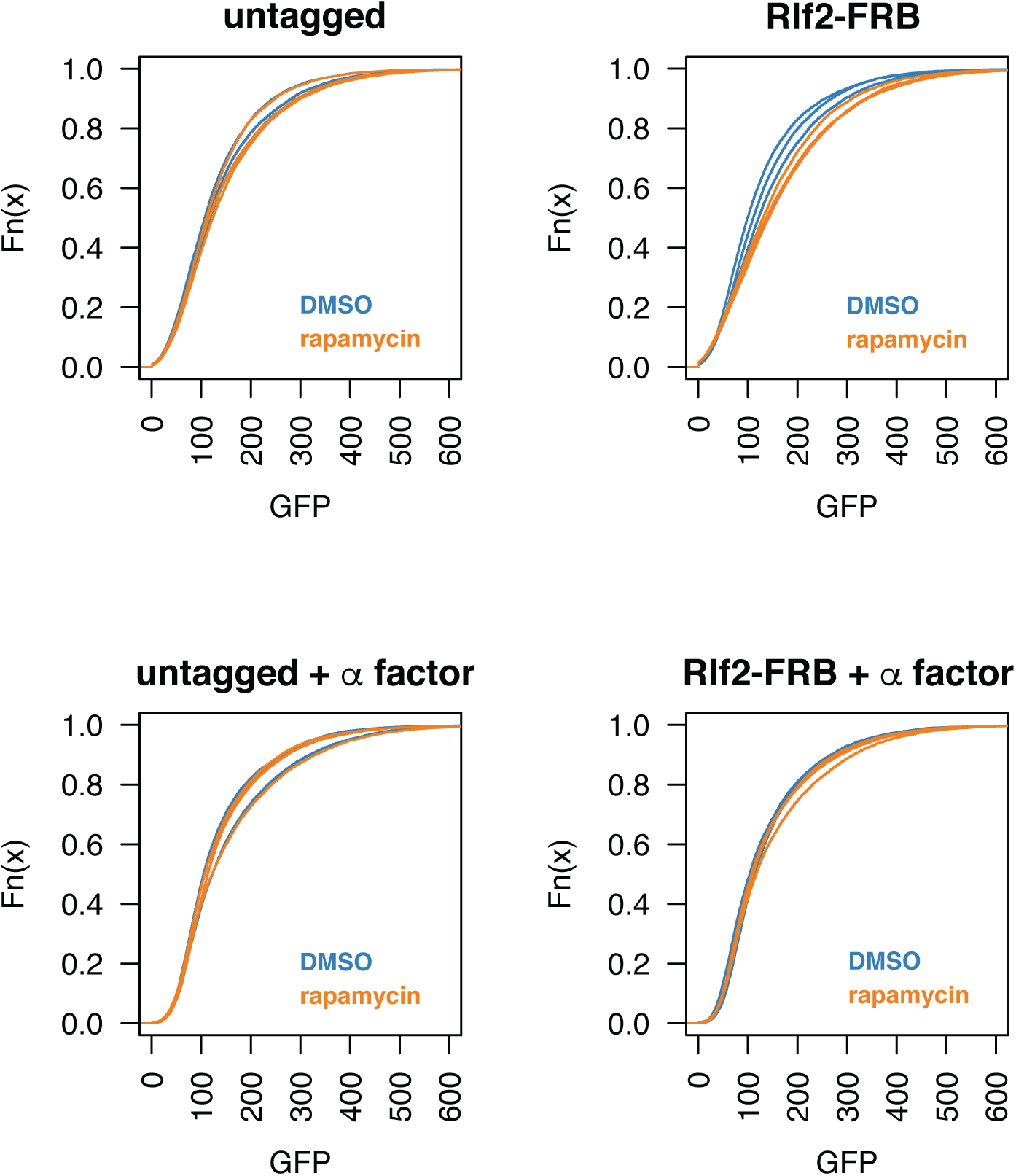
Cell-cycle arrest restores cryptic transcription suppression following depletions of Rlf2. Rlf2 was depleted using anchor-away by the addition of a C-terminal FRB tag and treatment with rapamycin, and transcription from a FLO8-sfGFP cryptic promoter was assessed by flow cytometry of 10,000 cells. Cells were grown in dextrose and were either asynchronous or arrested in G1 with a factor prior to rapamycin addition. Cultures lacking rapamycin were treated with DMSO as vehicle control. Assays were performed on three independent cultures and empirical cumulative distribution function plots of each are presented.

**Figure S7.**
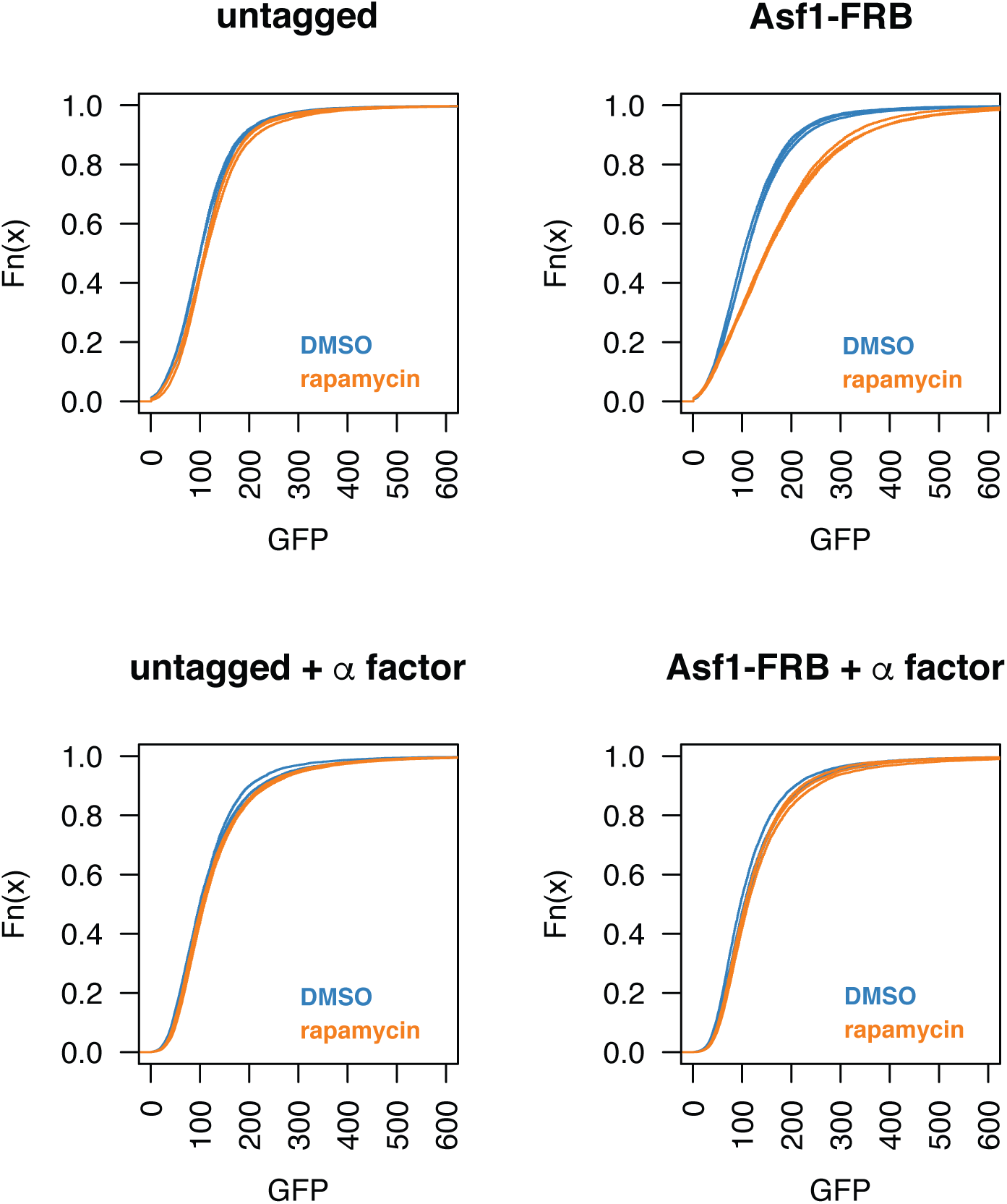
Cell cycle arrest rescues cryptic transcription defects due to depletion of Asf1. Asf1 was depleted using anchor-away by the addition of a C-terminal FRB tag and treatment with rapamycin (orange), and transcription from a *FLO8-sfGFP* cryptic promoter was assessed by flow cytometry of 10,000 cells. Cultures lacking rapamycin were treated with DMSO as vehicle control (blue). Cells were grown in dextrose and were either

**Figure S8.**
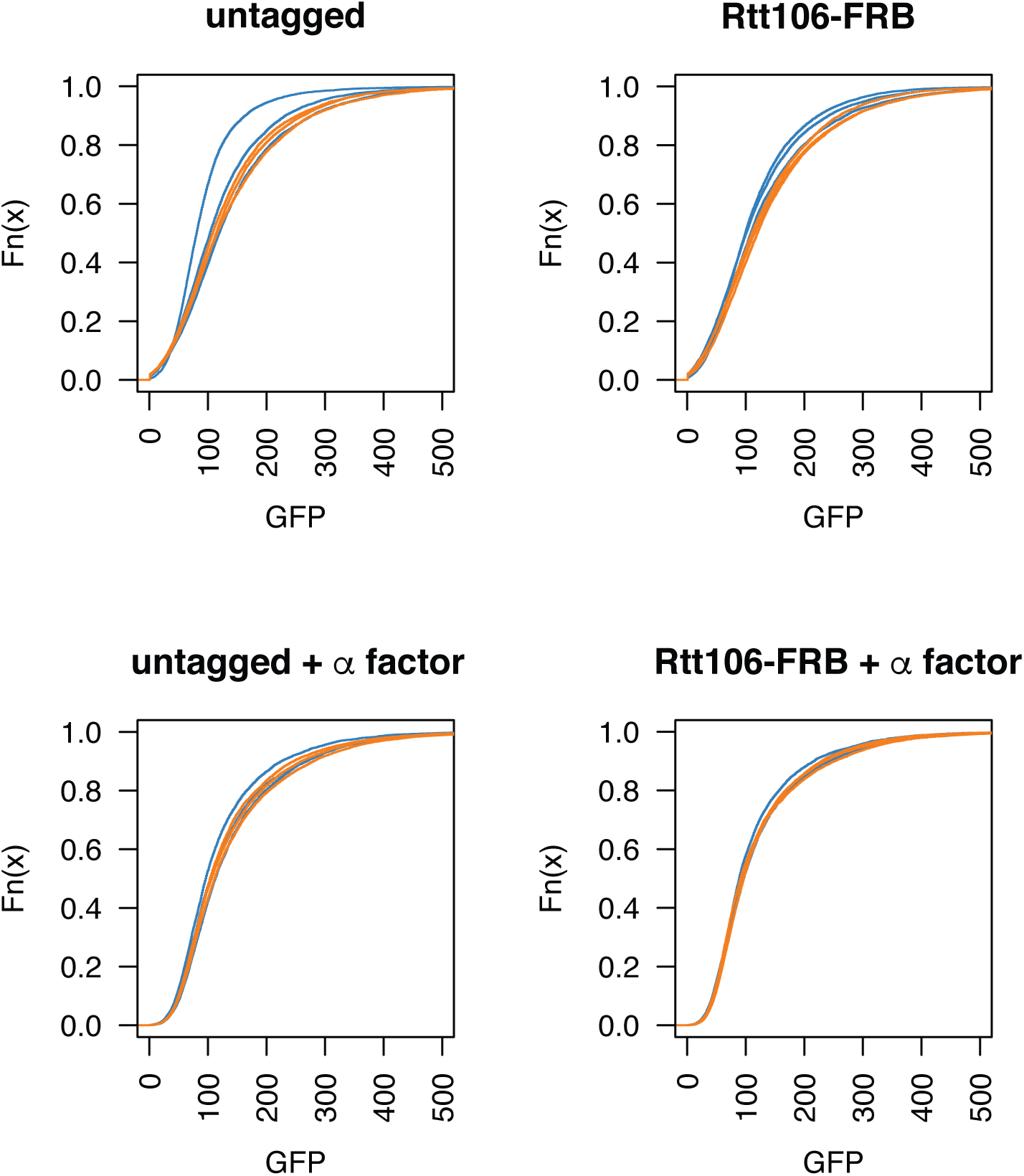
Cell cycle arrest rescues cryptic transcription defects due to depletion of Rtt106. Rtt106 was depleted using anchor-away by the addition of a C-terminal FRB tag and treatment with rapamycin (orange), and transcription from a *FLO8-sfGFP* cryptic promoter was assessed by flow cytometry of 10,000 cells. Cultures lacking rapamycin were treated with DMSO as vehicle control (blue). Cells were grown in dextrose and were either asynchronous or arrested in G1 with α factor prior to rapamycin/DMSO addition. Assays were performed on three independent cultures and empirical cumulative distribution function plots of each are presented.

**Figure S9.**
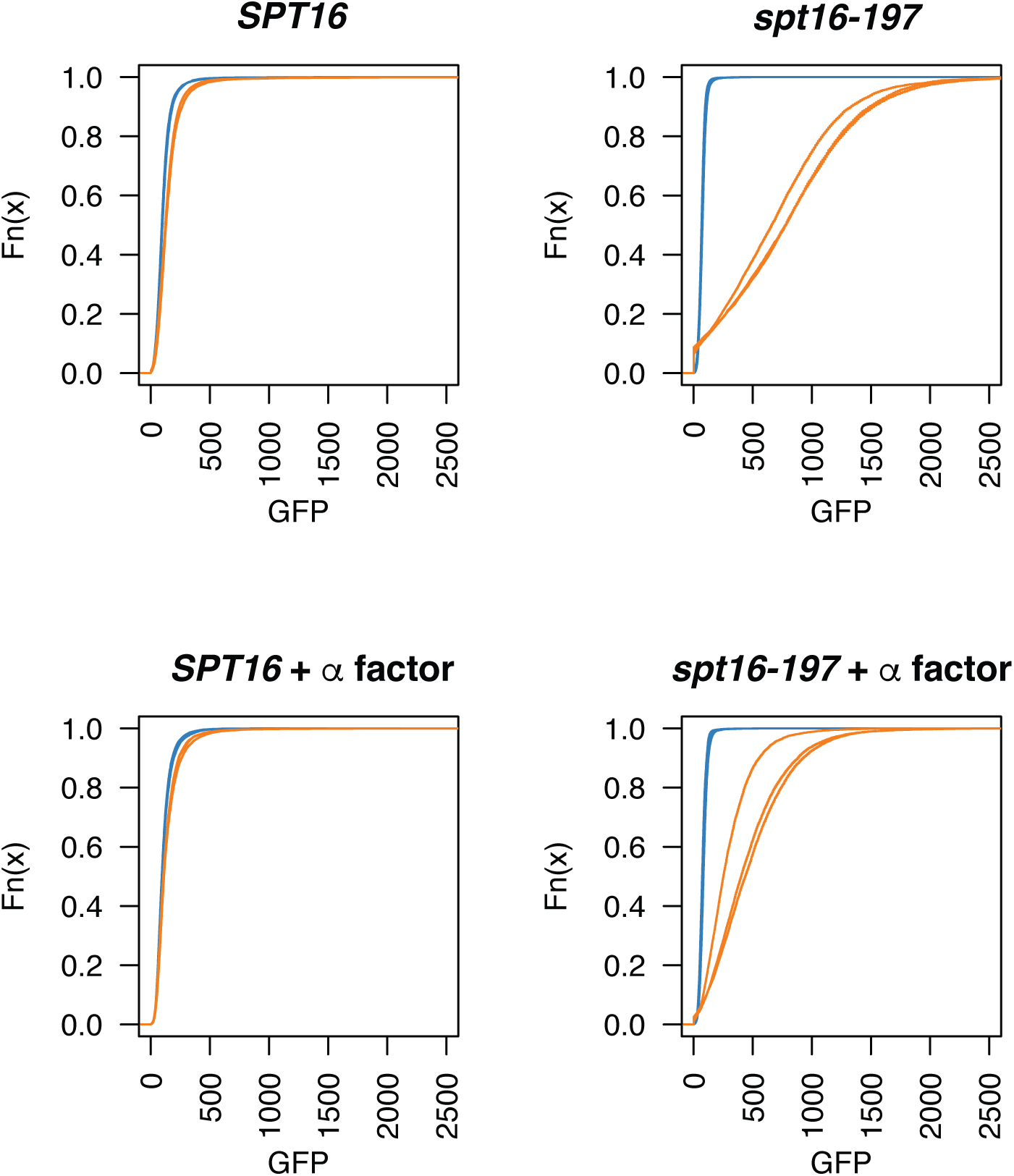
Cell cycle arrest rescues cryptic transcription defects in a *SPT16* mutant. The indicated strains were grown at 28°C (blue) or 37°C (orange) and transcription from a *FLO8*-sfGFP cryptic promoter was assessed by flow cytometry of 10,000 cells. Cells were either asynchronous or arrested in G1 with α factor prior to a shift to the non-permissive temperature. Assays were performed on three independent cultures and empirical cumulative distribution function plots of each are presented.

**Figure S10.**
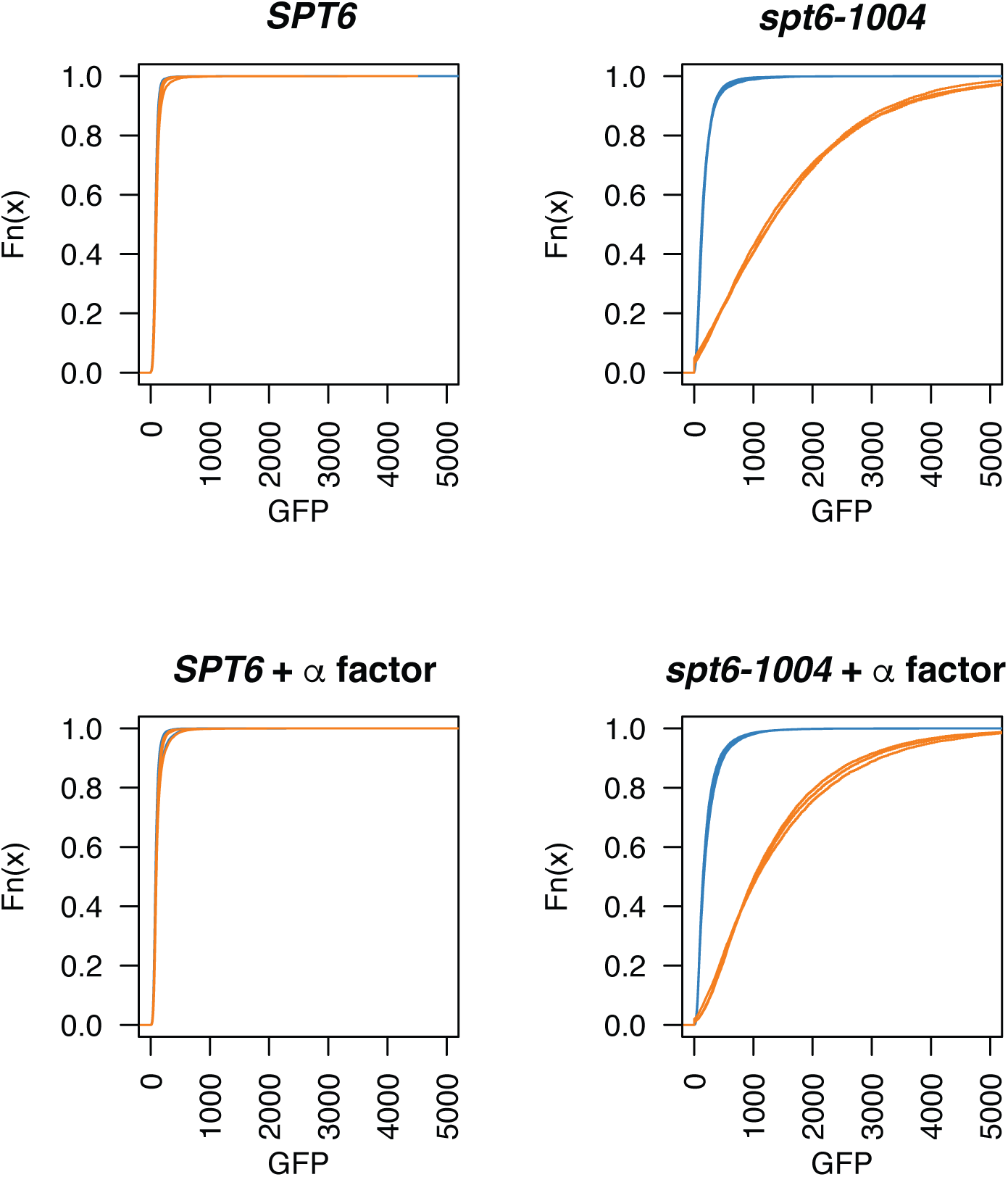
Cell cycle arrest rescues cryptic transcription defects in a *SPT6* mutant. The indicated strains were grown at 30°C (blue) or 37°C (orange) and transcription from a *FLO8*-sfGFP cryptic promoter was assessed by flow cytometry of 10,000 cells. Cells were either asynchronous or arrested in G1 with α factor prior to a shift to the non-permissive temperature. Assays were performed on three independent cultures and empirical cumulative distribution function plots of each are presented.

